# Rapid Assembly and Screening of Multivalent Immune Cell-Redirecting Therapies for Leukemia

**DOI:** 10.1101/2020.05.08.082628

**Authors:** Priscilla Do, Lacey A Perdue, Andrew Chyong, Rae Hunter, Jodi Dougan, Curtis J Henry, Christopher C Porter, Erik C Dreaden

**Affiliations:** Coulter Department of Biomedical Engineering, Georgia Institute of Technology and Emory University; Department of Pediatrics, Emory School of Medicine; Winship Cancer Institute of Emory University; Aflac Cancer and Blood Disorders Center, Children’s Healthcare of Atlanta and Emory School of Medicine; Petit Institute for Bioengineering and Bioscience, Georgia Institute of Technology

**Keywords:** nanotechnology, multivalency, leukemia, drug screening

## Abstract

Therapies that bind with immune cells and redirect their cytotoxic activity towards diseased cells represent a promising and versatile approach to immunotherapy with applications in cancer, lupus, and other diseases; traditional methods for discovering these therapies, however, are often time-intensive and lack the throughput of related target-based discovery approaches. Inspired by the observation that the cytokine, IL-12, can enhance antileukemic activity of the clinically approved T cell redirecting therapy, blinatumomab, here we describe the structure and assembly of a chimeric immune cell-redirecting agent which redirects the lytic activity of primary human T cells towards leukemic B cells and simultaneously co-targets the delivery of T cell-stimulating IL-12. We further describe a novel method for the parallel assembly of compositionally diverse libraries of these bi-specific T cell engaging cytokines (BiTEokines) and their high-throughput phenotypic screening, requiring just days for hit identification and the analysis of structure-function relationships. Using this approach, we identified CD19 × CD3 × IL12 compounds that exhibit *ex vivo* lytic activity comparable to current FDA-approved therapies for leukemia and correlated drug treatment with specific cell-cell contact, cytokine delivery, and leukemia cell lysis. Given the modular nature of these multivalent compounds and their rapid assembly/screening, we anticipate facile extension of this therapeutic approach to a wide range of immune cells, diseased cells, and soluble protein combinations in the future.

## INTRODUCTION

Immune cell redirection (ICR) is a powerful and versatile therapeutic approach in which the cytotoxic activity of endogenous immune cells is redirected towards diseased cells via simultaneous, drug-induced cell binding. This strategy has demonstrated therapeutic benefit in preclinical models of cancer,^1 2^ HIV,^3, 4^ lupus,^5^ and other diseases; however, only one such drug with an Fc-independent mechanism-of-action is currently approved for clinical use in the US: the bispecific antibody, blinatumomab, which redirects T cell killing towards leukemic B cells. Given the ability of ICR therapies to co-opt a wide range of cell types (e.g. T cells, NK cells,^6^ and macrophages^7^) against both cell-surface and intracellular targets,^8^ enthusiasm for future drug development is high with dozens of drug candidates at or in clinical-stage development.^2^

In addition to their diversity of application, ICR immunotherapies can also vary widely in their composition and mode of delivery. They encompass nanoparticle^9-11^, bispecific IgG,^12, 13^ scFv fusion,^14^ and mRNA^15^ constructs, as well as vectors based on oncolytic viruses^16^ and engineered cells.^17^ While a majority of ICR therapies in clinical testing are produced using traditional genetic engineering techniques, one challenge to their discovery and development is the relatively low-throughput manner in which drug candidates can be investigated and the relatively high dependency of drug action on the affinity of individual cell-binding domains. Fusion protein engineering methods that rely on conventional plasmid vectors^18^ or *de novo* protein design^19^ often require months for expression and purification prior to screening, and the effects of associated modifications on subsequent protein affinity can be challenging to predict.^20^ Moreover, while response rates to blinatumomab are often impressive, remissions are not always durable.^21^ Methods to both accelerate the discovery and improve the potency of ICR therapies are therefore urgently needed.

Recently, we identified IL-12 as a key mediator of the immune response to leukemia cells in mouse models of B cell acute lymphoblastic leukemia (ALL), including that recombinant IL-12 therapy alone could improve T cell activation, immunologic memory, and overall survival in mouse models of the disease.^22^ Based on these findings, we hypothesized that the activity of ICR therapies targeting T cells and leukemic B cells may be improved by concurrent delivery of IL-12, particularly if the two agents were tethered to one another in order to improve the typically poor circulation of IL-12 that limits its therapeutic potential.^23^

To examine this hypothesis, here we describe a method for the rapid assembly and screening of multivalent ICR drug candidates that redirect the lytic activity of T cells towards leukemic B cells and simultaneously co-deliver T cell-stimulating IL-12 to yield multifunctional therapies which we term, bispecific T cell-engaging cytokines (**BiTEokines**). Using this discovery platform, we show that cytokine co-delivery can dramatically alter the antileukemic activity of ICR immunotherapies and that the generation and screening of diverse libraries of BiTEokine candidates can be achieved in a matter of days, rather than weeks or months, thereby greatly accelerating the process of hit identification. Future extensions of this approach could enable the rapid identification of drug compounds with activity against cancer, autoimmune diseases, or pathogen infections and, given its modular nature, could be extended to a wide range of immune cells, diseased cells, and soluble proteins combinations in the future.

## RESULTS AND DISCUSSION

### IL-12 enhances bispecific T cell engager activity

We^22^ and others previously found that IL-12 can improve cancer immune elimination via enhanced CD8^+^ T cell proliferation,^24^ cytotoxicity,^17, 25^ survival,^17^ and T cell receptor (TCR) signaling.^26, 27^ As clinically approved T cell engager therapies are thought to act primarily on a subset of CD8^+^ T cells,^28, 29^ we posited that IL-12 may improve the lytic activity of blinatumomab which bi-specifically targets T cell CD3 and leukemic B cell CD19. Following prolonged co-culture of primary CD8^+^ T cells with CD19^+^ NALM-6 leukemia cells, we observed significant improvement in target leukemia cell lysis in the presence of IL-12, as measured by flow cytometry, (**Figure 1a-c**) as well as an associated increase in T cell proliferation (**Figure 1d**) and T cell activation (**Figure 1e**), as measured by dye dilution and interferon gamma (IFNγ) secretion, respectively. Together, these data demonstrate that IL-12 can improve the performance of T cell redirecting therapies in an *ex vivo* assay that is heavily relied upon to prioritize ICR drug candidates^30, 31^ and that these effects are attributable, in part, to cytokine-enhanced T cell proliferation and activation.

**Figure 1.**
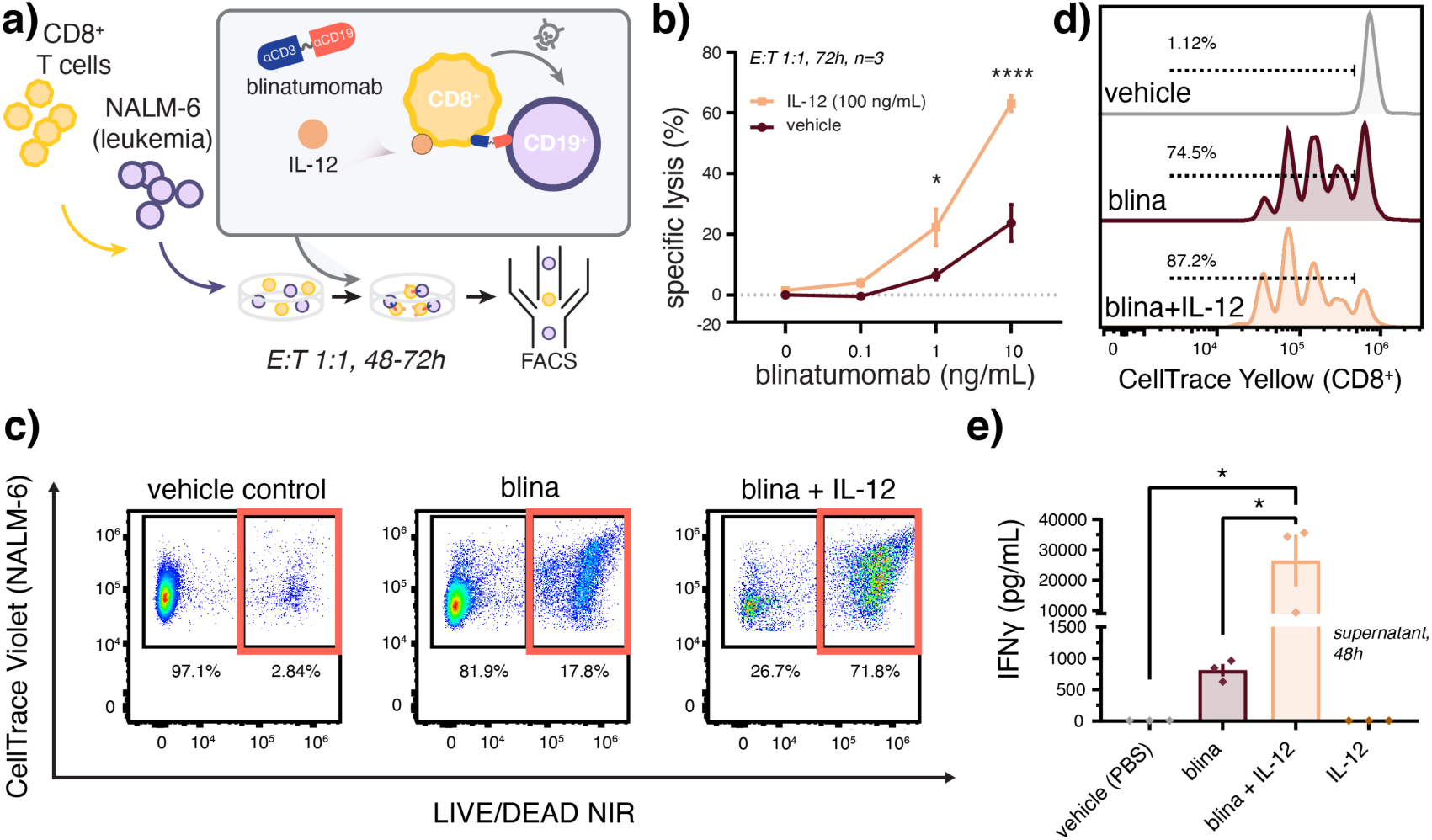
Recombinant IL-12 enhances activity of the bispecific T cell engager therapy, blinatumomab. **a)** Schematic of assay conditions for the co-culture of primary human CD8^+^ T cells with CD19^+^ NALM-6 leukemia cells. Blinatumomab-induced **(b**,**c)** lysis of NALM-6 leukemia cells and **(d)** T cell proliferation enhanced by co-incubation with IL-12 as measured by flow cytometry. **e)** Blinatumomab and IL-12 synergize to enhance T cell activation as measured by ELISA of IFNγ secretion into co-culture supernatants. Data in (c,d) report representative dot plots and dye-dilution histograms, respectively. Co-cultures in (c-e) were treated by blina (7 ng/mL) with or without IL-12 (3.5 ng/mL) in comparison to PBS vehicle over (c,d) 72 h or (e) 48 h. Values report (b) mean±SEM (n=3 donors) as analyzed by the mixed-effects model with correction for multiple comparisons and (e) mean±SEM (n=3 donors) as analyzed by one-way ANOVA with Tukey’s correction for multiple comparisons. See Supporting Information for associate gating strategies. *p<0.05, ****p<0.0001.

This marked effect of IL-12 on blinatumomab activity is significant in that other T cell mitogens such as IL-2 have been previously combined with blinatumomab with relatively little impact on lytic activity.^14, 30^ While the synergy observed here may be unique to IL-12, such differential effects may arise due to the fact that typical lysis assays are performed over much shorter durations (*e.g.* 4 h) and that the effects of IL-12 result, *in part*, due to cytokine-induced T cell proliferation (**Figure 1d**). Interestingly however, T cell expansion observed in the presence of both drugs was insufficient to account for the large change in both target cell lysis and T cell activation (via IFNγ secretion), thus future studies focusing on the role of IL-12 in modulating the selective expansion, differentiation,^32^ or activation CD8^+^ T cells in the presence of blinatumomab are warranted.

Also, while not investigated further in this work, the observation of synergy between blinatumomab and recombinant IL-12 is significant in that the former is currently approved to treat relapsed/refractory and minimal residual disease positive (MRD^+^) B-ALL in both adults in children. And while IL-12 therapies have not advanced to phase III trials due to poor circulation and toxicity, several novel IL-12 drug candidates currently under investigation may benefit from combination with blinatumomab including adenoviral,^33^ plasmid,^34^ mRNA,^35^ and affinity-targeted^24, 25^ IL-12.

### Design and rapid screening of CD19 × CD3 × IL12 BiTEokines

Having shown that IL-12 potentiates the antileukemic activity of T cell-redirecting immunotherapy, we next devised a drug architecture that (i) directs the lytic activity of T cells towards leukemic B cells, (ii) simultaneously co-delivers T cell-stimulating IL-12, and (iii) features a modular design amenable to combinatorial assembly of test compound libraries (**Figure 2**). We based the core scaffold of these structures on magnetic iron oxide nanoparticles due to their track-record of clinical use,^36, 37^ ability to accommodate a wide range of IgG antibodies via Fc-protein G affinity, and rapid purification via magnetic field sedimentation. Antibody clones were selected due to their prior clinical testing as CAR-T cell constructs (CD19, SJ25-C1)^38^ or antibody-drug conjugates (CD3, UCHT1),^39^ and their comparable IgG1-protein G affinity.

**Figure 2.**
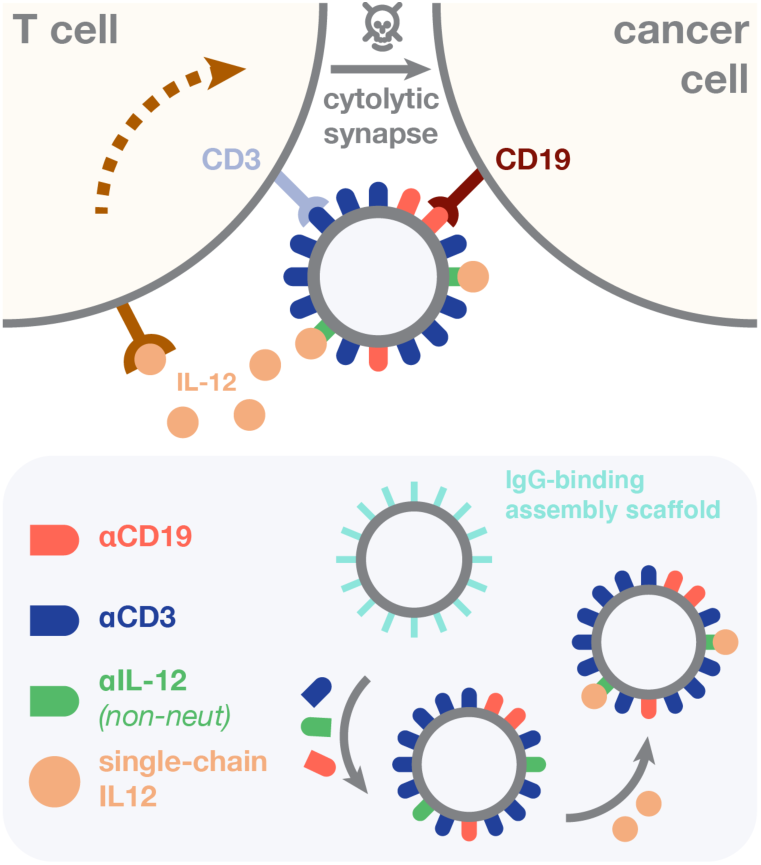
Structure and assembly of bispecific T cell engaging cytokines (BiTEokines). Schematic of drug-induced synapse formation between T cells and leukemic B cells, as well as synapse-targeted delivery of the cytokine, IL-12. Inset illustrates the modular and rapid self-assembly of CD19 × CD3 × IL12 BiTEokines via addition of human IgG to protein G-conjugated iron oxide nanoparticles and subsequent cytokine complexation.

To assemble BiTEokine test compound libraries, we dispensed equivalent numbers of magnetic nanoparticles into individual wells of a standard 96 well plate, each containing varying cocktails of fluorochrome-labeled antibodies directed against human CD19, CD3ε, (non-neutralizing) IL-12, or isotype control (**Figure 3a, S1**). After incubation, magnetic field-induced sedimentation, and analysis of antibody abundance using a standard fluorescence plate reader, we then added varying volumes of a novel single-chain variant of IL-12 (scIL-12), followed by incubation, purification, and sterile filtration. Using this methodology, BiTEokine libraries were prepared and characterized over the course of just 8 to 9 hours.

**Figure 3.**
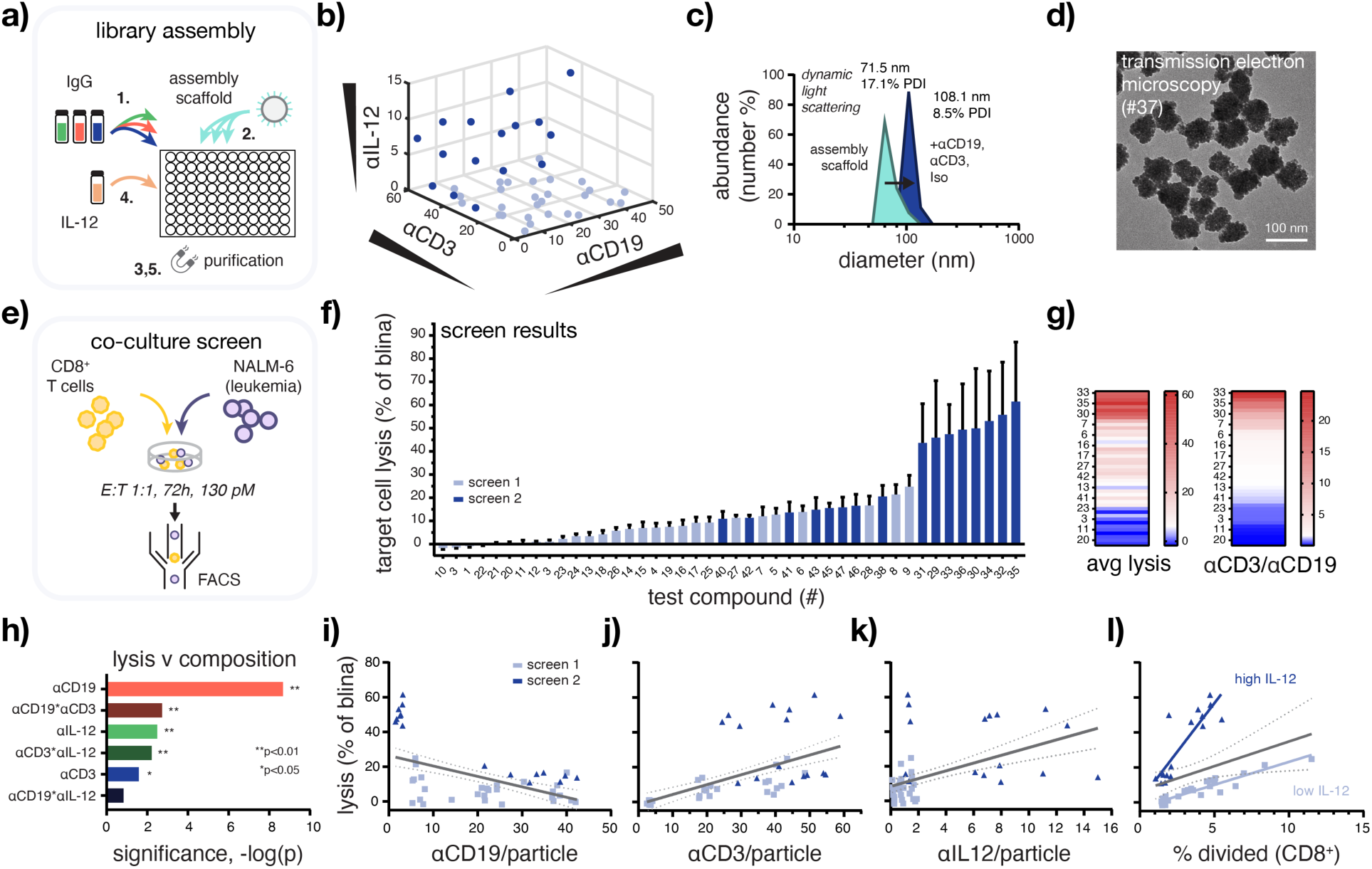
High-throughput assembly and screening enables rapid identification of BiTEokines that induce efficient leukemia cell lysis. **a)** Schematic of test compound library assembly. **b)** Structural diversity of the BiTEokine library and **(c**,**d)** representative test compound size/morphology as measured by antibody fluorescence intensity, dynamic light scattering, and transmission electron microscopy, respectively. **e)** Schematic of co-culture assay conditions and **(f)** parallel screening results rank-ordered by drug-induced lysis of CD19^+^ NALM-6 leukemia cells by primary human CD8^+^ T cells. **g)** Heatmaps illustrating concordance between target cell lysis the ratio of αCD3 to αCD19 per BiTEokine and **h)** significance of antibody components to measured lysis values as calculated by least squares modeling. Structure-function relationships reporting NALM-6 leukemia cell lysis versus **(i-k)** antibody abundance per particles and **(l)** T cell division. Values in (f) represent mean±SEM of 2-3 T cell donors, each analyzed in duplicate. Lines in (i-l) report linear regressions with 95% confidence intervals. *p<0.05, **p<0.01.

In total, we synthesized 47 unique BiTEokine test compounds which varied widely in antibody composition, achieving a consistent, and near theoretical maximum, total coverage of 134±15 IgG per particle (αCD19: 1.5±0.8 to 42±4; αCD3: 2.5±0.5 to 59±3; αIL12 0.12±0.08 to 15±0.7; **Figure 3b, S2**). Dynamic light scattering measurements indicated high stability of the subsequent test compounds in buffer, with hydrodynamic size increasing from 71.5 nm to 108.1 nm upon antibody surface-assembly (**Figure 3c**) with no appreciable change in particle morphology as measured by transmission electron microscopy (**Figure 3d**). We note that such size increases (ca. 36.5 nm) correspond closely to what one would expect following addition of a single monolayer of IgG1 (10-12 nm hydrodynamic size)^40^ about these particles. Together these data demonstrate that BiTEokine test compound libraries can be assembled rapidly, in parallel with a wide range of structural diversity.

With a test compound library in-hand, we next screened the lytic activity of BiTEokines following incubation with co-cultures of primary human CD8^+^ T cells and CD19^+^ NALM-6 leukemia cells and analysis by flow cytometry (**Figure 3e,f**). As anticipated, BiTEokine test compounds varied widely in their corresponding lytic potential with median activity just 11% (n=3) that of blinatumomab’s. Top screening hits, in contrast, exhibited lytic activity closely comparable to blinatumomab (#035: 62±26%). Interestingly, the top five performing BiTEokines displayed only a small number of B cell-targeting antibodies per particle (2±1 to 3.2±0.3) and an abundance for CD3 antibodies (26±1 to 59±3), thus largely limiting the potential for interaction with multiple B cells. Based on our prior studies in leukemia-bearing mice,^22^ we anticipate further therapeutic benefits from the delivery of IL-12 *in vivo*, due to its actions alone and when cross-exposed to antigens from lysed target cells.

Another attractive feature of this combinatorial screening approach is its ability to rapidly shed light on structure-function relationships unique to these novel multivalent drug architectures. For example, the bispecific antibody, blinatumomab, is well-known to target T and leukemic B cells with low and high affinity, respectively.^41^ Least squares regression modeling of screening data indicated significant contributions from all BiTEokine antibody components; however surprisingly, here we observed that high αCD3/αCD19 ratio was closely associated with favorable lytic activity (**Figure 3g,h**). Further, we found that αCD19 abundance was a negative predictor of lytic activity and, conversely, that αCD3, αIL12, and T cell division (% divided) were positive correlates of leukemia cell lysis. Interestingly, the impact of IL-12 on drug activity appeared to bifurcate depending on is relative abundance on BiTEokines with low amounts of cytokine (approx. 2.3 ng/mL) inducing high T cell division but low leukemia cell lysis, and high IL-12 (approx. 270 ng/mL) inducing less rapid T cell division and high target cell lysis (**Figure 3i-l**). While future studies will be required to attribute the origin of these differential effects from IL-12, we speculate that (i) the novel ability of multivalent BiTEokines to induce TCR clustering^42^ may allow them to redirect the activity of CD8^+^ T cell subsets outside of those typically acted upon by bispecific antibodies and (ii) that IL-12 concentration-dependent CD8^+^ T cell differentiation^32^ may enrich for T cell subsets with differential dependency on co-stimulation for drug-induced lysis (*e.g.* memory precursor or short-lived effector cells).

### Activity and specificity of BiTEokines

After identifying hits from the BiTEokine activity screen, we sought to confirm that these compounds specifically targeted T and leukemic B cells via flow cytometry. As anticipated, we observed particle abundance-dependent increases in the labeling of T cells with CD3 antibodies and B cells with CD19 antibodies, but no apparent αIL12-dependent cell specificity from BiTEokine test compounds (**Figure 4a**). In addition, we further confirmed that BiTEokine lytic activity arose from precise combinations of antibodies, rather than non-specific antibody interactions, using compounds either fully or partially conjugated with isotype control antibody to approximately equivalent total amounts of antibody. While isotype control BiTEokines elicited only basal levels of activity (1.9% lysis), we observed >17-fold increases in lytic responses from lead compounds #35 and 37 (**Figure 4b**). Together, these data correlate cell-specific binding by BiTEokines with the lysis of leukemic B cells.

**Figure 4.**
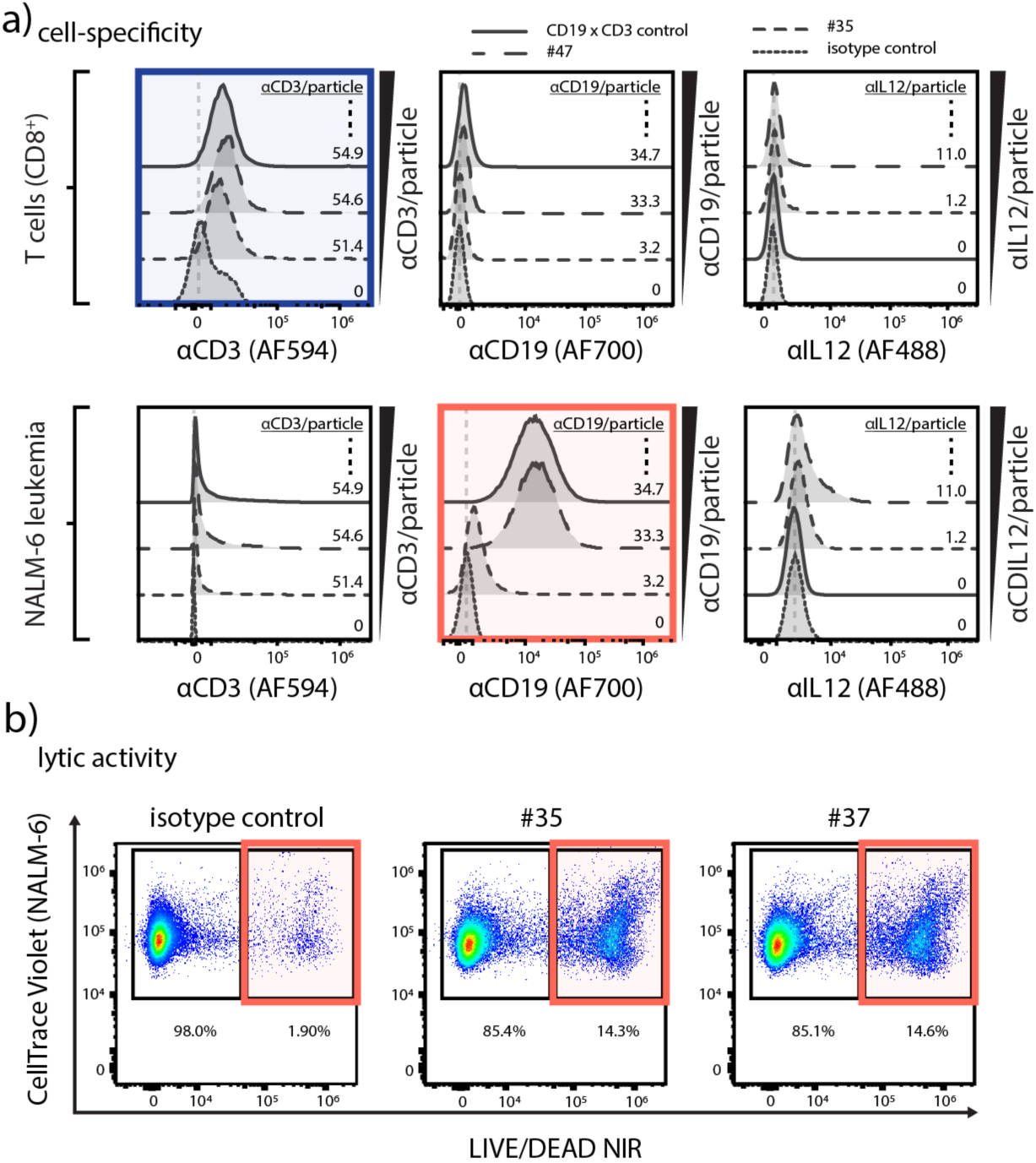
CD19 × CD3 × IL12 BiTEokines bind specifically and induce efficient leukemia cell lysis. **a)** Cell fluorescence from various BiTEokine antibodies observed in co-cultures gated on (top) T cells or (bottom) NALM-6 leukemia cell as measured by flow cytometry. *Top left:* CD8^+^ T cells exhibit CD3 antibody fluorescence that increases in intensity with relative abundance on BiTEokines. *Bottom middle:* CD19^+^ leukemia cells exhibit CD19 antibody fluorescence that increases in intensity with relative abundance on BiTEokines. **b)** Representative dot plots of BiTEokine-induced NALM-6 leukemia cell killing in comparison to isotype control-conjugated particles. Experimental conditions in (a,b) are noted in Fig 3.

To further characterize the role of BiTEokines in inducing leukemia cell lysis, we performed imaging flow cytometry on co-cultures of primary human CD8^+^ T cells and CD19^+^ NALM-6 leukemia cells treated with fluorescently labeled hit compound #35 from the prior activity screen, as well as blinatumomab. Gating on doublets of T and B cells, we observed similar patterns of LAMP-1 (CD107a) positive vesicle accumulation, indicative of lytic granules and lysosomes, in both blinatumomab- and BiTEokine-treated co-cultures (**Figure 5**). Strikingly, we observed distinct accumulation of BiTEokines at the interface between T and leukemic B cells, further correlating BiTEokine treatment with cell-cell contact with leukemia cell lysis.

**Figure 5.**
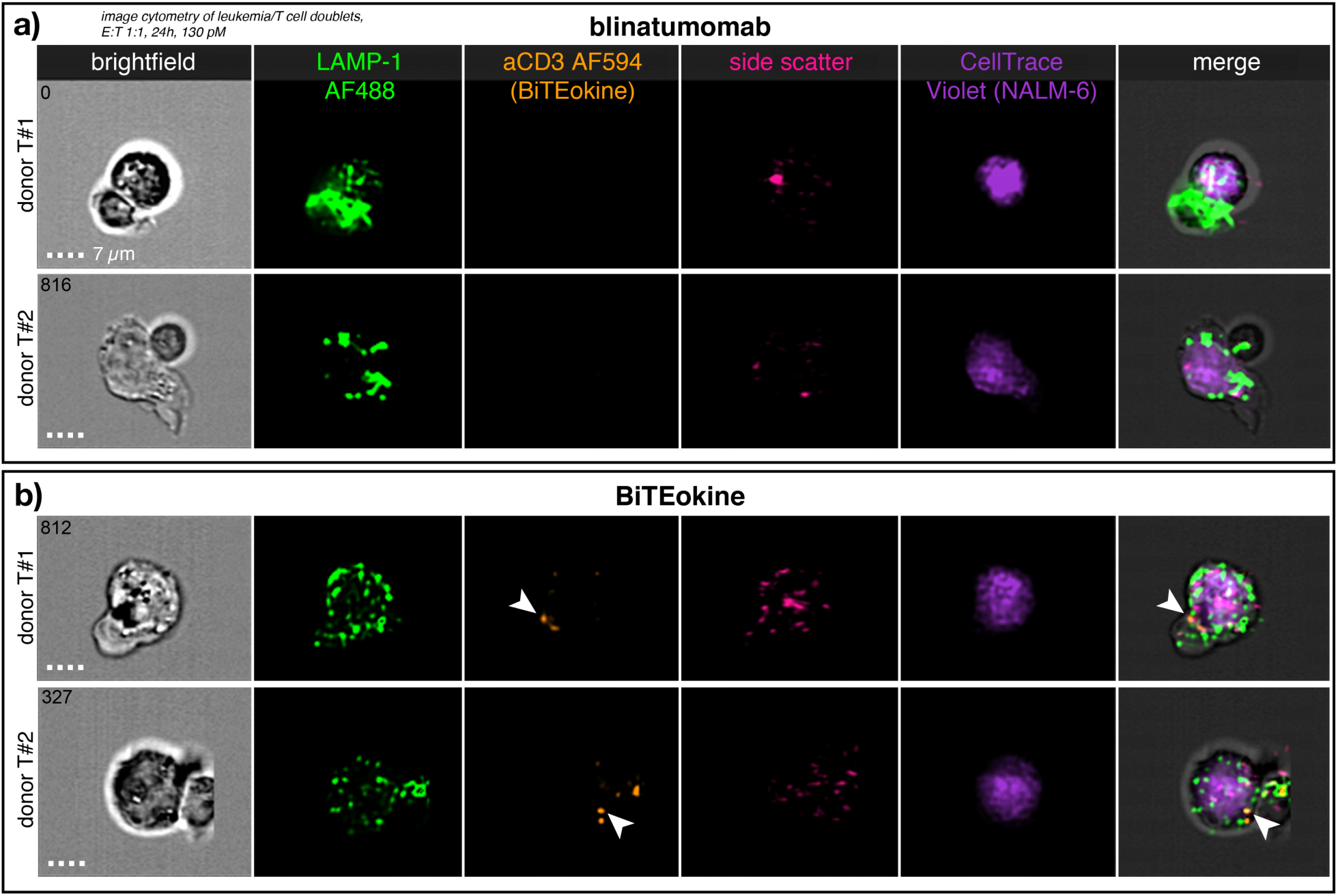
BiTEokines localize at the interface between primary human T cells and NALM-6 leukemic B cells. Imaging flow cytometry of CD8^+^ T cells co-cultured with NALM-6 leukemia cells at 1:1 E:T ratio and treated **(a)** blinatumomab or **(b)** CD19 × CD3 × IL12 BiTEokines (#35) for 24 h at equimolar concentrations (130 pM). (a,b) report images from two different T cell donors. Arrowheads indicate localization of BiTEokines at the T-B cell interface. Scale bar is 7*µ*m.

## CONCLUSIONS

Here, we describe a methodology for the rapid discovery of immune cell-redirecting therapies which are combinatorially self-assembled from recombinant proteins and magnetic nanoparticles. Motivated by the antileukemic activity of IL-12, both alone and in combination with the T cell engager therapy, blinatumomab, we devised a modular and convergently assembled drug structure that redirects the lytic activity of T cells towards leukemic B cells and simultaneously co-targets the delivery of T cells-stimulating IL-12. We show that compositionally diverse libraries of these CD19 × CD3 × IL12 BiTEokines can be assembled and screened over the course of just days – rather than months that are typically required using traditional recombinant techniques – enabling rapid hit identification and the delineation of important structure-function relationships. Using this approach, we identified BiTEokine hit compounds which exhibit *ex vivo* lytic activity comparable to current FDA-approved therapies for leukemia. Detailed analysis of BiTEokine activity strongly correlated drug treatment with specific cell-cell contact, IL-12 delivery, and leukemia cell lysis. These results are particularly promising given that we anticipate additional *in vivo* therapeutic benefit from IL-12, due to immune memory resulting from cross-exposure of cytokine with antigens from lysed target cells. Although future lead optimization may be required in order to maximize *in vivo* activity from these compounds, these studies demonstrate that optimal antibody composition and density can be rapidly determined using this approach; such information could be used to inform the synthesis of structurally analogous antibody-conjugated liposomes, virus-like particles (VLPs),^46, 47^ and self-assembled protein cages^48^ for subsequent translation. Future studies investigating the impact of core scaffold size and targeted cytokine neutralization, rather than delivery, may also lead to further improvements in BiTEokine activity or an expansion of disease targets, respectively. While a limited number of promising synthetic ICR agents have been previously described,^9-11^ these studies are the first – to our knowledge – to present a method for the discovery and screening-based optimization of this promising class of immunotherapy. Given the rapid and modular nature of the approach presented here, we also anticipate facile extension to a wide range of immune cells, diseased cells, and soluble protein combinations in the future.

## METHODS

### Primary cells and cell lines

De-identified, normal donor blood samples were obtained from ZenBio (Durham, NC). PBMCs were isolated from buffy coats by Ficoll density gradient centrifugation. CD8^+^ T cells were isolated from PBMCs by negative selection using EasySep (Human CD8+ T cell Isolation Kit, Stemcell), assessed for ≥ 90% purity by flow cytometry, and cryopreserved. Primary cells and the NALM-6 cell line (gifted from Dr. Lia Gore, University of Colorado) were cultured in RPMI (10% FBS, 100 U/mL penicillin, 100 μg/mL streptomycin). Hek-Blue IL-12 reporter cells were obtained from Invivogen and cultured in DMEM (10% FBS, 50 U/mL penicillin, 100 μg/mL streptomycin, 100 μg/mL Normocin, 1x HEK-Blue selection). All cells were cultured at 37 °C in a 5% CO_2_ humidified atmosphere and tested regularly for mycoplasma.

### Therapeutic antibodies and recombinant proteins

Fluorochrome-conjugated IgG antibodies were purchased from Biolegend: anti-CD3 (UCHT1), anti-CD19 (SJ25-C1), anti-IL-12 (C11.5), and IgG1 isotype control (MOPC21C). Single chain human IL-12 was purchased from Invivogen. Blinatumomab (anti-hCD19-CD3) was obtained from. Protein concentrations were measured via UV optical absorption (Nanodrop, Thermo).

### BiTEokine synthesis and characterization

Magnetic nanoparticles (50-80nm) functionalized with protein G were obtained from Ocean Nanotech. Particles were validated for lot-specific sizing via DLS (DynaPro III, Wyatt). BiTEokine test compounds were prepared via addition of 1.65 × 10^11^ particles to antibody mixtures (PBS) for 10 min at room temperature with agitation at 200 rpm. Antibody amounts per well ranged from: 0.11 μg to 2.28 μg (αCD19), 1.25 μg to 24.31 μg (αCD3), 0.91 μg to 8.38 μg (αIL12), 1.76 μg to 11.27 μg (Isotype). Unbound antibodies were removed by via magnetic field-induced sedimentation (≤ 8 min) and washing twice with PBS. Previously obtained calibration curves for antibody-particle binding were used to establish conditions for compound library preparation. Intermediate compounds were then passed through 0.45*µ*m sterile cellulose acetate filters and 2 eq. of recombinant human single chain IL-12 (relative to αIL12 binding sites) was added to a subset of particles for 30 min at room temperature with agitation at 200 rpm. Purified test compounds were obtained after magnetic field-induced sedimentation (≥ 8 min) and washing twice with PBS. Antibody abundance on test compounds was determined from spillover-corrected fluorescence intensity and comparison to standard curves for each fluorochrome-conjugated antibody.

BiTEokine test compounds were imaged via transmission electron microscopy at 80 kV using a Hitachi HT-7700 instrument following sample application to formvar/carbon coated copper grids (400 mesh, Electron Microscopy Sciences) for 15 minutes and washing for 2 seconds with ultrapure water. Hydrodynamic size was measured via dynamic light scattering using a DynaPro III plate reader (Wyatt).

### Standard Flow Cytometry

Primary cells were stained for purity post-isolation using anti-CD45 (HI30, BD), anti-CD8 (RPA-T8, BD), and Near IR Live/dead stain (Invitrogen), then fixed with 4% formaldehyde (Thermo) and analyzed using a BD LSR II or a Cytek Aurora cytometer. Data were analyzed using FlowJo 10 software.

### Imaging Flow Cytometry

Cell multimers were fixed by gently adding 4% formaldehyde directly to cell co-cultures to a final concentration of 2% for 15 min at room temperature. Cells were then permeabilized with saponin (BD Perm/wash), stained with anti-LAMP1 AF488 (eBioH4A3, Thermo), washed 2x and resuspended in PBS, then analyzed using an ImageStream^x^ Mk II (Amnis) instrument with 60x magnification in extended depth-of-field mode. Data were analyzed using IDEAS software (Amnis).

### Cytotoxicity Assay

Target leukemia cells were stained with CFSE (Tonbo) or CellTrace Violet (Thermo) and T cells were left unstained or stained with CellTrace Yellow prior to co-culture. Target cells and T cells were co-cultured in 96 well u-bottom plates for up to 72 h. Count beads (Invitrogen) were added to each sample to determine absolute cell counts. After co-culture, cells were stained with Near IR Live/Dead for viability and fixed with 4% formaldehyde prior to flow cytometric analysis. Test compounds for which multiple donors or multiple replicates were not obtained, were excluded from analysis. Specific lysis was calculated using the equation: %Specific Lysis= 100 × (%Treated Sample [Violet+NIR LD+] – %IsoBeads[Violet+NIR LD+])/(100-%IsoBeads[Violet+NIR LD+]).

### Proliferation Assay

Proliferation of T cells was measured by dye dilution of CellTrace Yellow (Thermo). Gating was performed using FlowJo software and analysis of dye dilution data was performed using ModFit software (Verity) to determine proliferation index and percent divided.

### ELISA Assay

Human IFNγ was quantified via sandwich ELISA assay (430107, Biolegend) and per the manufacturer’s recommended conditions.

### Statistics and Software

Analyses were performed in Graphpad Prism, JMP Pro 14, and Modfit. Statistical comparisons were performed via one-way or two-way ANOVA with correction for multiple comparisons using Graphpad Prism. Analysis of BiTEokine variables contributing to optimal lytic activity was performed via standard least squares modeling in JMP Pro 14.

## ACKNOWLEDGEMENTS

This work was supported in part by the US Department of Defense (Idea Award CA180783), the AAI Careers in Immunology Fellowship Program, the National Institutes of Health Research Training Program in Immunoengineering (T32EB021962), the American Cancer Society (IRG-17-181-05), the Coulter Department of Biomedical Engineering, and the Aflac Cancer and Blood Disorders Center of Children’s Healthcare of Atlanta. We are also grateful for assistance from the Children’s Healthcare of Atlanta and Emory University’s Pediatric Integrated Cellular Imaging Core and Pediatric General Equipment & Specimen Processing Core, the Robert P. Apkarian Integrated Electron Microscopy Core, and the Emory Chemical Biology Discovery Center. The content here is solely the responsibility of the authors and does not necessarily represent the official views of the organizations acknowledged herein.

## AUTHOR CONTRIBUTIONS

P.D., C.J.H., C.C.P., and E.C.D. designed research; P.D., L.A.P., A.C., R.H., J.D., C.J.H., C.C.P., and E.C.D. performed research or analyzed data; P.D., L.A.P., R.H., C.J.H., C.C.P., and E.C.D. edited the manuscript, and C.J.H., C.C.P., and E.C.D. wrote the manuscript.

## COMPETING INTERESTS

The authors declare no competing interests.

## TABLE OF CONTENTS IMAGE

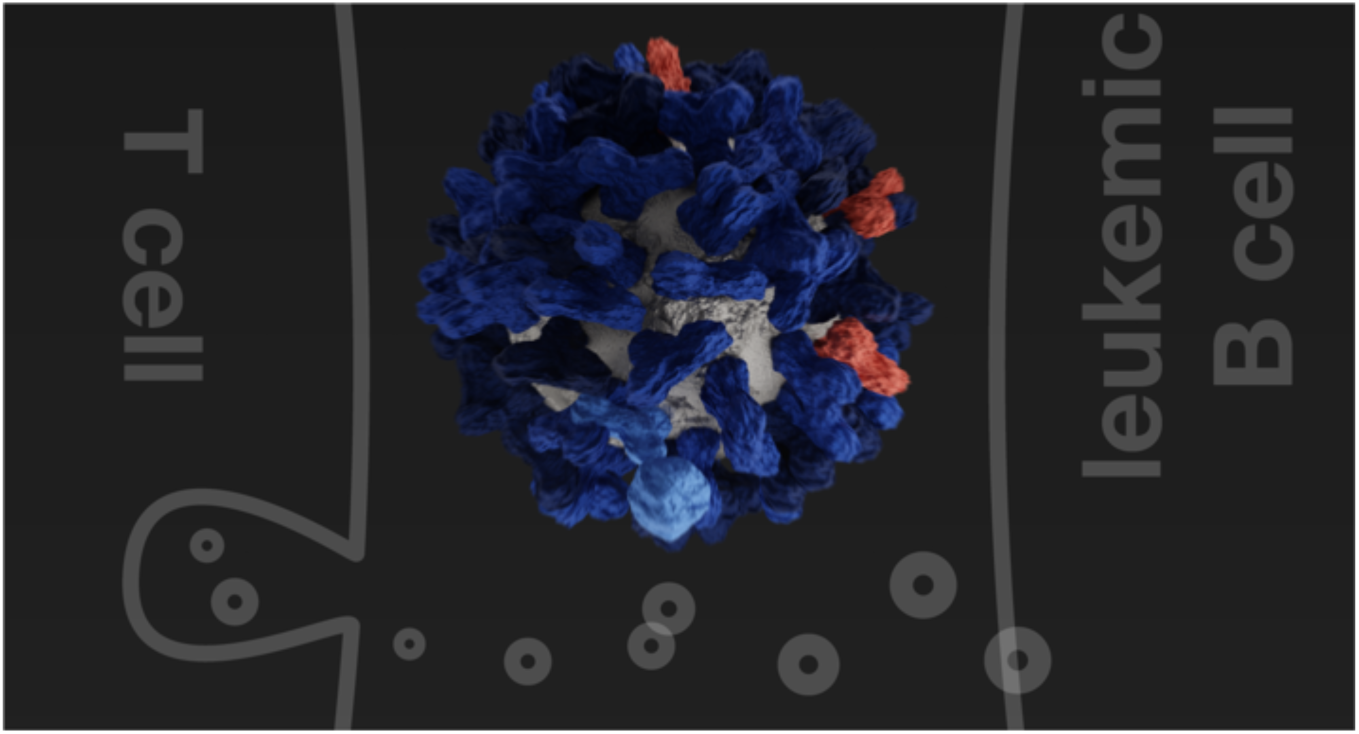

## Notes

### Competing Interest Statement

The authors have declared no competing interest.

## REFERENCES

1. Baeuerle, P. A.; Reinhardt, C., Bispecific T-Cell Engaging Antibodies for Cancer Therapy. Cancer Res. 2009, 69 (12), 4941–4944.

2. Goebeler, M.-E.; Bargou, R. C., T cell-engaging therapies — BiTEs and beyond. Nat. Rev. Clin. Oncol. 2020.

3. Bardhi, A.; Wu, Y.; Chen, W.; Li, W.; Zhu, Z.; Zheng, J. H.; Wong, H.; Jeng, E.; Jones, J.; Ochsenbauer, C.; Kappes, J. C.; Dimitrov, D. S.; Ying, T.; Goldstein, H., Potent In Vivo NK Cell-Mediated Elimination of HIV-1-Infected Cells Mobilized by a gp120-Bispecific and Hexavalent Broadly Neutralizing Fusion Protein. J. Virol. 2017, 91 (20), e00937–17.

4. Brozy, J.; Schlaepfer, E.; Mueller, C. K. S.; Rochat, M.-A.; Rampini, S. K.; Myburgh, R.; Raum, T.; Kufer, P.; Baeuerle, P. A.; Muenz, M.; Speck, R. F., Antiviral Activity of HIV gp120-Targeting Bispecific T Cell Engager Antibody Constructs. Journal of Virology 2018, 92 (14), e00491–18.

5. Schooten, W. v.; Juelke, K.; Iyer, S.; Buelow, B.; Buelow, R.; Volk, D., 293 A novel CD3/BCMA bispecific antibody selectively kills plasma cells in bone marrow of healthy individuals with improved safety. Lupus Sci. Med. 2019, 6 (Suppl 1), A213–A213.

6. Vallera, D. A.; Felices, M.; McElmurry, R.; McCullar, V.; Zhou, X.; Schmohl, J. U.; Zhang, B.; Lenvik, A. J.; Panoskaltsis-Mortari, A.; Verneris, M. R.; Tolar, J.; Cooley, S.; Weisdorf, D. J.; Blazar, B. R.; Miller, J. S., IL15 Trispecific Killer Engagers (TriKE) Make Natural Killer Cells Specific to CD33+ Targets While Also Inducing Persistence, In Vivo Expansion, and Enhanced Function. Clin. Cancer Res. 2016, 22 (14), 3440–3450.

7. Scott, E. M.; Jacobus, E. J.; Lyons, B.; Frost, S.; Freedman, J. D.; Dyer, A.; Khalique, H.; Taverner, W. K.; Carr, A.; Champion, B. R.; Fisher, K. D.; Seymour, L. W.; Duffy, M. R., Bi- and tri-valent T cell engagers deplete tumour-associated macrophages in cancer patient samples. J. Immunotherapy Cancer 2019, 7 (1), 320.

8. Liddy, N.; Bossi, G.; Adams, K. J.; Lissina, A.; Mahon, T. M.; Hassan, N. J.; Gavarret, J.; Bianchi, F. C.; Pumphrey, N. J.; Ladell, K.; Gostick, E.; Sewell, A. K.; Lissin, N. M.; Harwood, N. E.; Molloy, P. E.; Li, Y.; Cameron, B. J.; Sami, M.; Baston, E. E.; Todorov, P. T.; Paston, S. J.; Dennis, R. E.; Harper, J. V.; Dunn, S. M.; Ashfield, R.; Johnson, A.; McGrath, Y.; Plesa, G.; June, C. H.; Kalos, M.; Price, D. A.; Vuidepot, A.; Williams, D. D.; Sutton, D. H.; Jakobsen, B. K., Monoclonal TCR-redirected tumor cell killing. Nat. Med. 2012, 18 (6), 980–987.

9. Schütz, C.; Varela, J. C.; Perica, K.; Haupt, C.; Oelke, M.; Schneck, J. P., Antigen-specific T cell Redirectors: a nanoparticle based approach for redirecting T cells. Oncotarget 2016, 7 (42).

10. Yuan, H.; Jiang, W.; von Roemeling, C. A.; Qie, Y.; Liu, X.; Chen, Y.; Wang, Y.; Wharen, R. E.; Yun, K.; Bu, G.; Knutson, K. L.; Kim, B. Y. S., Multivalent bi-specific nanobioconjugate engager for targeted cancer immunotherapy. Nat. Nanotech. 2017, 12 (8), 763–769.

11. Vaidya, T.; Straubinger, R. M.; Ait-Oudhia, S., Development and Evaluation of Tri-Functional Immunoliposomes for the Treatment of HER2 Positive Breast Cancer. Pharm. Res. 2018, 35 (5), 95.

12. Staerz, U. D.; Bevan, M. J., Hybrid hybridoma producing a bispecific monoclonal antibody that can focus effector T-cell activity. Proc. Natl. Acad. Sci. U.S.A. 1986, 83 (5), 1453–1457.

13. Burris, H. A.; Giaccone, G.; Im, S.-A.; Bauer, T. M.; Oh, D.-Y.; Jones, S. F.; Nordstrom, J. L.; Li, H.; Carlin, D. A.; Baughman, J. E.; Lechleider, R. J.; Bang, Y.-J., Updated findings of a first-in-human, phase I study of margetuximab (M), an Fc-optimized chimeric monoclonal antibody (MAb), in patients (pts) with HER2-positive advanced solid tumors. J. Clin. Oncol. 2015, 33 (15_suppl), 523–523.

14. Löffler, A.; Kufer, P.; Lutterbüse, R.; Zettl, F.; Daniel, P. T.; Schwenkenbecher, J. M.; Riethmüller, G.; Dörken, B.; Bargou, R. C., A recombinant bispecific single-chain antibody, CD19□×□CD3, induces rapid and high lymphoma-directed cytotoxicity by unstimulated T lymphocytes. Blood 2000, 95 (6), 2098–2103.

15. Stadler, C. R.; Bähr-Mahmud, H.; Celik, L.; Hebich, B.; Roth, A. S.; Roth, R. P.; Karikó, K.; Türeci, Ö.; Sahin, U., Elimination of large tumors in mice by mRNA-encoded bispecific antibodies. Nat. Med. 2017, 23 (7), 815–817.

16. Speck, T.; Heidbuechel, J. P.; Veinalde, R.; Jaeger, D.; von Kalle, C.; Ball, C. R.; Ungerechts, G.; Engeland, C. E., Targeted BiTE expression by an oncolytic vector augments therapeutic efficacy against solid tumors. Clin. Cancer Res. 2018, clincanres.2651.2017.

17. Yeku, O. O.; Purdon, T. J.; Koneru, M.; Spriggs, D.; Brentjens, R. J., Armored CAR T cells enhance antitumor efficacy and overcome the tumor microenvironment. Sci. Rep. 2017, 7 (1), 10541.

18. Sampei, Z.; Haraya, K.; Tachibana, T.; Fukuzawa, T.; Shida-Kawazoe, M.; Gan, S. W.; Shimizu, Y.; Ruike, Y.; Feng, S.; Kuramochi, T.; Muraoka, M.; Kitazawa, T.; Kawabe, Y.; Igawa, T.; Hattori, K.; Nezu, J., Antibody engineering to generate SKY59, a long-acting anti-C5 recycling antibody. PLOS ONE 2018, 13 (12), e0209509.

19. Perkel, J. M., The computational protein designers. Nature 2019, 571, 585–587.

20. Zuch de Zafra, C. L.; Fajardo, F.; Zhong, W.; Bernett, M. J.; Muchhal, U. S.; Moore, G. L.; Stevens, J.; Case, R.; Pearson, J. T.; Liu, S.; McElroy, P. L.; Canon, J.; Desjarlais, J. R.; Coxon, A.; Balazs, M.; Nolan-Stevaux, O., Targeting Multiple Myeloma with AMG 424, a Novel Anti-CD38/CD3 Bispecific T-cell–recruiting Antibody Optimized for Cytotoxicity and Cytokine Release. Clin. Cancer Res. 2019, 25 (13), 3921–3933.

21. Kantarjian, H.; Stein, A.; Gökbuget, N.; Fielding, A. K.; Schuh, A. C.; Ribera, J.-M.; Wei, A.; Dombret, H.; Foà, R.; Bassan, R.; Arslan, Ö.; Sanz, M. A.; Bergeron, J.; Demirkan, F.; Lech-Maranda, E.; Rambaldi, A.; Thomas, X.; Horst, H.-A.; Brüggemann, M.; Klapper, W.; Wood, B. L.; Fleishman, A.; Nagorsen, D.; Holland, C.; Zimmerman, Z.; Topp, M. S., Blinatumomab versus Chemotherapy for Advanced Acute Lymphoblastic Leukemia. New Engl. J. Med. 2017, 376 (9), 836–847.

22. Rabe, J. L.; Gardner, L.; Hunter, R.; Fonseca, J. A.; Dougan, J.; Gearheart, C. M.; Leibowitz, M. S.; Lee-Miller, C.; Baturin, D.; Fosmire, S. P.; Zelasko, S. E.; Jones, C. L.; Slansky, J. E.; Rupji, M.; Dwivedi, B.; Henry, C. J.; Porter, C. C., IL-12 abrogates calcineurin-dependent immune evasion during leukemia progression. Cancer Res. 2019, canres.3800.2018.

23. Berraondo, P.; Etxeberria, I.; Ponz-Sarvise, M.; Melero, I., Revisiting Interleukin-12 as a Cancer Immunotherapy Agent. Clin Cancer Res 2018, 24 (12), 2716–2718.

24. Mansurov, A.; Ishihara, J.; Hosseinchi, P.; Potin, L.; Marchell, T. M.; Ishihara, A.; Williford, J.-M.; Alpar, A. T.; Raczy, M. M.; Gray, L. T.; Swartz, M. A.; Hubbell, J. A., Collagen-binding IL-12 enhances tumour inflammation and drives the complete remission of established immunologically cold mouse tumours. Nat. Biomed Eng. 2020.

25. Momin, N.; Mehta, N. K.; Bennett, N. R.; Ma, L.; Palmeri, J. R.; Chinn, M. M.; Lutz, E. A.; Kang, B.; Irvine, D. J.; Spranger, S.; Wittrup, K. D., Anchoring of intratumorally administered cytokines to collagen safely potentiates systemic cancer immunotherapy. Sci. Transl. Med. 2019, 11 (498), eaaw2614.

26. Braun, M.; Ress, M. L.; Yoo, Y.-E.; Scholz, C. J.; Eyrich, M.; Schlegel, P. G.; Wölfl, M., IL12-mediated sensitizing of T-cell receptor-dependent and -independent tumor cell killing. OncoImmunology 2016, 5 (7), e1188245.

27. Goplen, N. P.; Saxena, V.; Knudson, K. M.; Schrum, A. G.; Gil, D.; Daniels, M. A.; Zamoyska, R.; Teixeiro, E., IL-12 Signals through the TCR To Support CD8 Innate Immune Responses. J. Immunol. 2016, 197 (6), 2434–2443.

28. Aigner, M.; Feulner, J.; Schaffer, S.; Kischel, R.; Kufer, P.; Schneider, K.; Henn, A.; Rattel, B.; Friedrich, M.; Baeuerle, P. A.; Mackensen, A.; Krause, S. W., T lymphocytes can be effectively recruited for ex vivo and in vivo lysis of AML blasts by a novel CD33/CD3-bispecific BiTE antibody construct. Leukemia 2013, 27 (5), 1107–1115.

29. Chames, P.; Baty, D., Bispecific antibodies for cancer therapy. mAbs 2009, 1 (6), 539–547.

30. Dreier, T.; Lorenczewski, G.; Brandl, C.; Hoffmann, P.; Syring, U.; Hanakam, F.; Kufer, P.; Riethmuller, G.; Bargou, R.; Baeuerle, P. A., Extremely potent, rapid and costimulation-independent cytotoxic T-cell response against lymphoma cells catalyzed by a single-chain bispecific antibody. Int. J. Cancer 2002, 100 (6), 690–697.

31. Hoffmann, P.; Hofmeister, R.; Brischwein, K.; Brandl, C.; Crommer, S.; Bargou, R.; Itin, C.; Prang, N.; Baeuerle, P. A., Serial killing of tumor cells by cytotoxic T cells redirected with a CD19-/CD3-bispecific single-chain antibody construct. Int. J. Cancer 2005, 115 (1), 98–104.

32. Joshi, N. S.; Cui, W.; Chandele, A.; Lee, H. K.; Urso, D. R.; Hagman, J.; Gapin, L.; Kaech, S. M., Inflammation Directs Memory Precursor and Short-Lived Effector CD8+ T Cell Fates via the Graded Expression of T-bet Transcription Factor. Immunity 2007, 27 (2), 281–295.

33. Barrett, J. A.; Cai, H.; Miao, J.; Khare, P. D.; Gonzalez, P.; Dalsing-Hernandez, J.; Sharma, G.; Chan, T.; Cooper, L. J. N.; Lebel, F., Regulated intratumoral expression of IL-12 using a RheoSwitch Therapeutic System® (RTS®) gene switch as gene therapy for the treatment of glioma. Cancer Gene Ther. 2018, 25 (5), 106–116.

34. Algazi, A.; Bhatia, S.; Agarwala, S.; Molina, M.; Lewis, K.; Faries, M.; Fong, L.; Levine, L. P.; Franco, M.; Oglesby, A.; Ballesteros-Merino, C.; Bifulco, C. B.; Fox, B. A.; Bannavong, D.; Talia, R.; Browning, E.; Le, M. H.; Pierce, R. H.; Gargosky, S.; Tsai, K. K.; Twitty, C.; Daud, A. I., Intratumoral delivery of tavokinogene telseplasmid yields systemic immune responses in metastatic melanoma patients. Ann. Oncol. 2020, 31 (4), 532–540.

35. Luheshi, N.; Hewitt, S.; Garcon, F.; Burke, S.; Watkins, A.; Arnold, K.; Zielinski, J.; Martin, P.; Sulikowski, M.; Bagnall, C.; Lapointe, J.-M.; Moody, G.; Si, H.; Morehouse, C.; Wilkinson, R. W.; Herbst, R.; Frederick, J., Abstract 5017: MEDI1191, a novel IL-12 mRNA therapy for intratumoral injection to promote TH1 transformation of the patient tumor microenvironment. Cancer Res. 2019, 79 (13 Supplement), 5017–5017.

36. Lu, M.; Cohen, M. H.; Rieves, D.; Pazdur, R., FDA report: Ferumoxytol for intravenous iron therapy in adult patients with chronic kidney disease. Am. J. Hematol. 2010, 85 (5), 315–319.

37. McCarthy, J. R.; Weissleder, R., Multifunctional magnetic nanoparticles for targeted imaging and therapy. Adv. Drug Del. Rev. 2008, 60 (11), 1241–1251.

38. Ramos, C. A.; Savoldo, B.; Dotti, G., CD19-CAR Trials. Cancer J. 2014, 20 (2), 112–118.

39. Frankel, A. E.; Woo, J. H.; Ahn, C.; Foss, F. M.; Duvic, M.; Neville, P. H.; Neville, D. M., Resimmune, an anti-CD3ε recombinant immunotoxin, induces durable remissions in patients with cutaneous T-cell lymphoma. Haematologica 2015, 100 (6), 794–800.

40. Hawe, A.; Hulse, W. L.; Jiskoot, W.; Forbes, R. T., Taylor Dispersion Analysis Compared to Dynamic Light Scattering for the Size Analysis of Therapeutic Peptides and Proteins and Their Aggregates. Pharm. Res. 2011, 28 (9), 2302–2310.

41. Zimmerman, Z.; Maniar, T.; Nagorsen, D., Unleashing the clinical power of T cells: CD19/CD3 bi-specific T cell engager (BiTE®) antibody construct blinatumomab as a potential therapy. Int. Immunol. 2014, 27 (1), 31–37.

42. Minguet, S.; Swamy, M.; Alarcón, B.; Luescher, I. F.; Schamel, W. W. A., Full Activation of the T Cell Receptor Requires Both Clustering and Conformational Changes at CD3. Immunity 2007, 26 (1), 43–54.

43. Giavridis, T.; van der Stegen, S. J. C.; Eyquem, J.; Hamieh, M.; Piersigilli, A.; Sadelain, M., CAR T cell–induced cytokine release syndrome is mediated by macrophages and abated by IL-1 blockade. Nat. Med. 2018, 24 (6), 731–738.

44. Norelli, M.; Camisa, B.; Barbiera, G.; Falcone, L.; Purevdorj, A.; Genua, M.; Sanvito, F.; Ponzoni, M.; Doglioni, C.; Cristofori, P.; Traversari, C.; Bordignon, C.; Ciceri, F.; Ostuni, R.; Bonini, C.; Casucci, M.; Bondanza, A., Monocyte-derived IL-1 and IL-6 are differentially required for cytokine-release syndrome and neurotoxicity due to CAR T cells. Nat. Med. 2018, 24 (6), 739–748.

45. Bluemel, C.; Hausmann, S.; Fluhr, P.; Sriskandarajah, M.; Stallcup, W. B.; Baeuerle, P. A.; Kufer, P., Epitope distance to the target cell membrane and antigen size determine the potency of T cell-mediated lysis by BiTE antibodies specific for a large melanoma surface antigen. Cancer Immunol. Immunother. 2010, 59 (8), 1197–1209.

46. Wang, Q.; Lin, T.; Tang, L.; Johnson, J. E.; Finn, M. G., Icosahedral Virus Particles as Addressable Nanoscale Building Blocks. Angew. Chemie Int. Ed. 2002, 41 (3), 459–462.

47. Gupta, S. S.; Kuzelka, J.; Singh, P.; Lewis, W. G.; Manchester, M.; Finn, M. G., Accelerated Bioorthogonal Conjugation: A Practical Method for the Ligation of Diverse Functional Molecules to a Polyvalent Virus Scaffold. Bioconjugate Chem. 2005, 16 (6), 1572–1579.

48. Bale, J. B.; Gonen, S.; Liu, Y.; Sheffler, W.; Ellis, D.; Thomas, C.; Cascio, D.; Yeates, T. O.; Gonen, T.; King, N. P.; Baker, D., Accurate design of megadalton-scale two-component icosahedral protein complexes. Science 2016, 353 (6297), 389–394.

