## Supplemental Information for "Rapid Assembly and Screening of Multivalent Immune Cell-Redirecting Therapies for Leukemia"

### SUPPORTING INFORMATION

*Gating Strategy*

Figure 1. Gating is presented as (b) time>singlets>FS/SS above debris>live/dead NIR+CFSE+ (NALM-6), (c) time>cells excluding count beads by far red+>cells excluding debris by FS/SS> CellTrace Violet+> live/dead NIR+), (d) time>cells excluding count beads by far red+>cells excluding debris by FS/SS> NIR-/CellTrace Violet- live T cells> CellTrace Yellow+ dividing cells.

Figure 4. Gating is presented as (a) time>cells excluding count beads by far red+>cells excluding debris by FS/SS> live cells by live/dead NIR-> CellTrace Yellow+/CellTrace Violet- (T cells) or CellTrace Yellow-/CellTrace Violet+ (Nalm6 cells) > anti-CD3 AF594, anti-CD19 AF700, or anti-IL-12 AF488 and (b) time>cells excluding count beads by far red+>cells excluding debris by FS/SS> CellTrace Violet+> live/dead NIR+.

Figure 5. Gating is presented as (a,b) low aspect ratio> CellTrace Violet+>LAMP-1 AF488+.

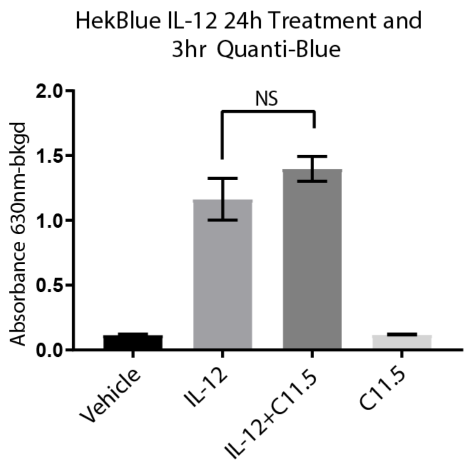


**Figure S1. Anti-IL-12 monoclonal antibody, clone C11.5, is non-neutralizing.** Soluble IL-12 activity (100 ng/mL, 24 h) in the presence or absence of αIL-12 as measured using IL-12 reporter cells.

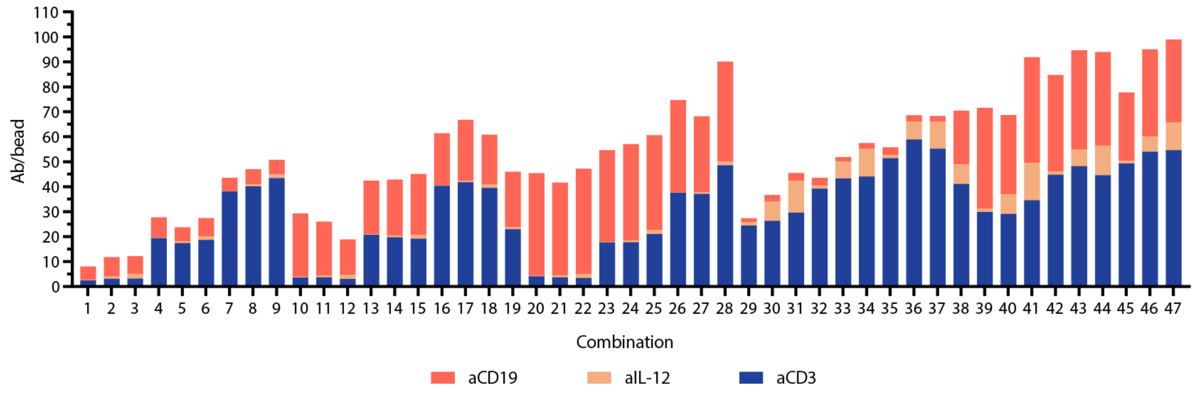


**Figure S2. Composition of the CD19 x CD3 x IL12 BiTEokine test compound library as measured by antibody fluorescence intensity.**
